# A Lipidomics Approach to Determine the Role of Lipids and its Crosstalk with Autophagy in Lung Cancer Metastasis

**DOI:** 10.1101/2024.01.01.573842

**Authors:** Simone C. da Silva Rosa, Javad Alizadeh, Rui Vitorino, Arun Surendran, Amir Ravandi, Biniam Kidane, Saeid Ghavami

## Abstract

Non-small cell lung cancer (NSCLC) is among the most malignant tumors with high propensity for metastasis and is the leading cause of cancer-related death globally. Most patients present with regional and distant metastasis, associated with poor prognosis. Lipids may play an essential role in either activating or inhibiting detachment-induced apoptosis (anoikis), where the latter is a crucial mechanism to prevent metastasis, and it may have a cross-talk with autophagy. Autophagy has been shown to be induced in various human cancer metastasis, modulating tumor cell motility and invasion, cancer cell differentiation, resistance to anoikis, and epithelial to mesenchymal transition. Hence, it may play a crucial role in the transition of benign to malignant phenotypes, the core of metastasis initiation. Here, we provide a method we have established in our laboratory for detecting of lipids in attached and detached non-small lung cancer cells and show how to analyze lipidomics data to find its correlation with autophagy-related pathways.

**Graphical abstract:** 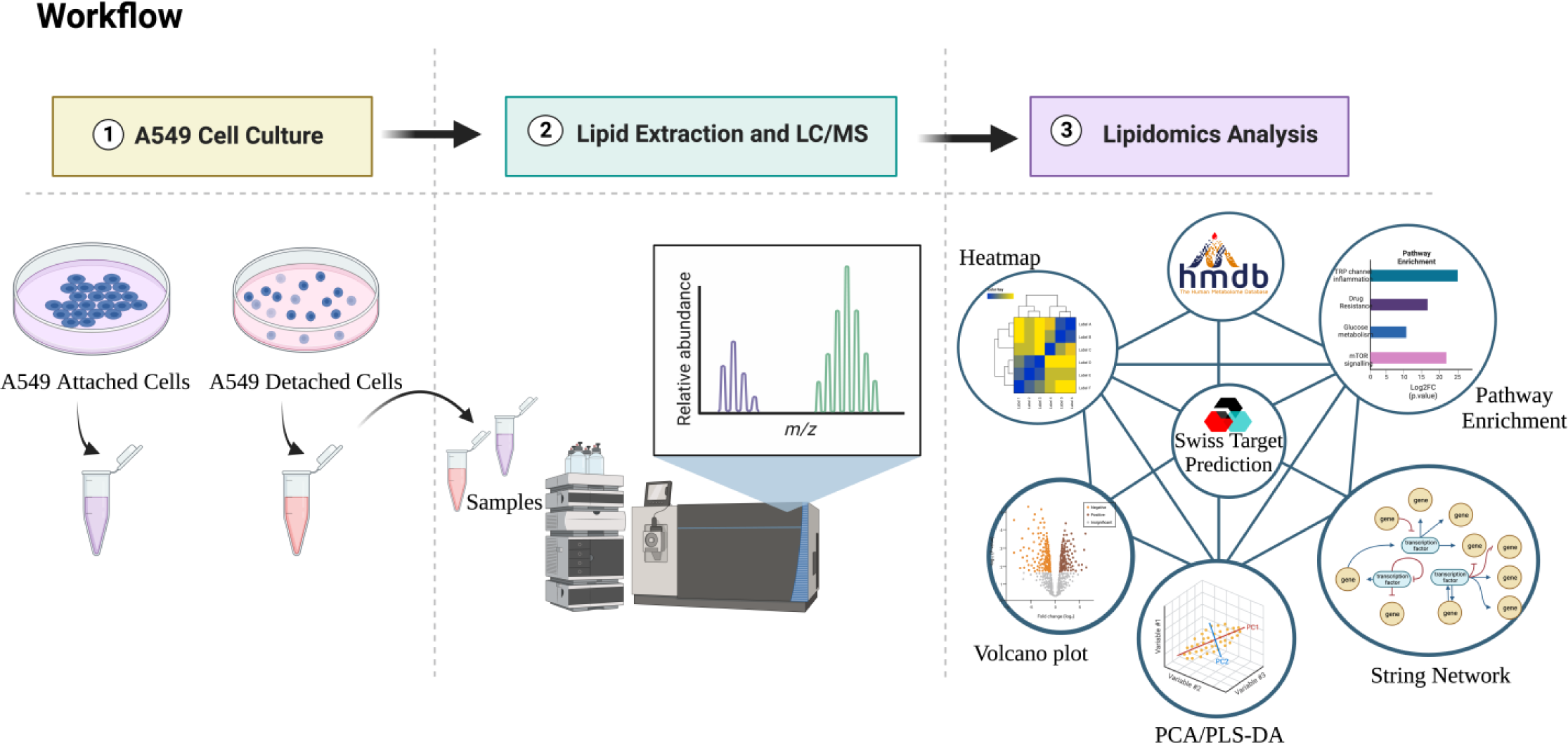

Workflow of A549 attached and detached cell culture. Cells are collected, lipids are extracted and subjected to LC/MS processing. Next, lipidomics resulting data is analyzed through various steps involving utilization of Metaboanlyst, HDMB, Swiss Target Prediction, and String.

## 1. Introduction

Non-small cell lung cancer (NSCLC) is among the most malignant tumors with high propensity for metastasis and is the leading cause of cancer-related death globally^1^. The NSCLC accounts for approximately 85% of all lung cancers worldwide, where the most common subtype is adenocarcinoma^2,3^. Many patients present with regional and distant metastasis^4^, associated with poor prognosis^5^. However, much yet must be elucidated on the precise mechanisms of NSCLC progression; hence, we need a deeper understanding of this issue to effectively identify effective diagnostic biomarkers and novel therapeutic targets.

Autophagy is an intracellular recycling mechanism that targets cytoplasmatic components to lysosomes for degradation and has been implicated in cancer^6^. After the autophagic breakdown of encapsulated lipids, proteins, carbohydrates or nucleic acids is completed, the product macromolecules return to the cytoplasm to support homeostatic cellular processes such as metabolism^7^. Although questionable, autophagy’s role in cancer has been reported as a dual-edged sword, promoting both tumor suppression and proliferation, depending on the developmental stage of the tumor, regulating mitochondrial function and stress survival^7–9^. Studies have shown that autophagy is induced in various human cancer metastasis, modulating tumor cell motility and invasion, cancer cell differentiation, resistance to anoikis, and epithelial to mesenchymal transition^10,11^. As a result, autophagy may play an essential role in the transition of benign to malignant phenotypes, the core of metastasis initiation^12^.

Cancer cells have advanced mechanisms to escape cell death, hijacking the programmed cell death, the natural barrier in cancer progression initiated by apoptosis mechanisms, and additional programmed cell death types, such as autophagy^13,14^. Lipid metabolism is another critical player in the regulation of cancer cell fate. The growing tumor cells heavily depend on lipid metabolic plasticity and reprogramming to sustain metabolic adaptations. As cancer cells develop a malignant phenotype, lipid metabolism changes, presenting increased lipid synthesis, storage and/or uptake, ultimately contributing to cancer progression^15^.

Over the recent years, various studies have demonstrated the complex interplay between lipid metabolism and autophagy in regulating cell death and survival^16–18^. Lipids play an important role in stress-induced autophagic membrane structure formation^19^. Moreover, the association between lipid metabolism and autophagy is well demonstrated through commonly shared regulators such as AMPK and mTORC signaling molecules, among others^20–22^. Lipid metabolism has been suggested to modulate autophagy during cancer progression^23^; hence, therapeutic strategies targeting lipid metabolism and autophagy may potentially enhance treatment effectiveness, eventually improving cancer patient survival^14^.

Guo et al (2013) showed that autophagy suppression in lung tumor maintains lipid homeostasis^24^. Previously, we have shown that Simvastatin, a widely used cholesterol-lowering agent, could further sensitize cancer cells (lung, breast, and brain) to temozolomide-induced cell death via autophagy inhibition^25,26^. Following that first observation, we became particularly interested in understanding the role of lipids and lung cancer metastasis and its crosstalk with autophagy. Therefore, using a lipidomics approach in our studies, we hope to elucidate the molecular mechanisms of NSCLC metastasis to determine one or more reliable early metastasis biomarkers to improve outcomes for NSCLC patients. Mass spectrometry is a common method choice for qualitative and quantitative lipidomics analysis, allowing the identification of many individual lipids^27^. The recent advance in this field culminated in the rapid abundance of big data in cancer omics, which, combined with computational resources and bioinformatics, accelerate our fundamental understanding of cancer biology to translational advancements^28^.

Here, we propose that lipids may play an important role in changing detachment-induced apoptosis (anoikis), where the latter is a crucial mechanism to prevent metastasis, and that it may have a cross-talk with autophagy. To further delineate the predominant lipids involved in metastasis, we cultured an adenocarcinoma human cell line (A549) in both attached and detached conditions, mimicking primary and metastatic tumor environments, and subjected these cells to a lipidomics screening through mass spectrometry. We then quantified the lipid abundance and performed an enrichment analysis of possible altered pathways resulting from predominantly altered lipids in both conditions.

In the materials and methods section, we will list and describe respectively the following steps:

1. A549 cell culture
2. Lipid extraction and LC/MS
3. Lipidomics analysis

## 2 Materials

### 2.1 A549 cell culture

1. Lung adenocarcinoma cells (A549; ATCC).
2. Adherent and low adherence plates culture plates.
3. Cell culture media: Dulbecco’s Modified Eagle’s Medium high glucose (4 mg/ml) (DMEM), 10 % Fetal Bovine Albumin (FBS), 1% Penicilin-Streptomycin, 1% insulin transferrin selenium (ITS).

### 2.2 Lipid chromatography and mass spectrometry^29^

#### 2.2.1 Lipid extraction

1. Phosphate Buffer Saline 1x (PBS), 1L: 800ml of distilled water, 8g of NaCl, 1.44g of Na2HPO4, 0.2g of KCl, 0.24g of KH2PO4, pH 7.4 or 7.2, top up with additional water until reaches 1L.
2. The lipids were quantified by comparing the lipid peak areas to those of the class-specific internal standards (ISTD). ISTD were added to the samples before the lipid extraction. The use of an ISTD is essential since they help to account for unavoidable variations in sample processing including suppression of ionization due to matrix interference, and manual discrepancies in lipid extraction. The choice of an ISTD for a particular lipid class depends upon the structural similarity and likeness in MS/MS fragmentation pattern of the species present in that lipid class.
3. In this study, 30 μL of ISTD in chloroform/methanol (1:1, v/v) were added to 200 μL of chloroform/methanol (2:1, v/v) along with 100 μL of cell homogenate. The details of internal samples and their concentration is provided below in **Table 1**.

**Table 1.** Lipid Internal Standards and Quantification used in this study.

| Lipid Class/Subclass | Internal Standard | $\mu\text{M}/30\ \mu\text{l}$ |
| --- | --- | --- |
| SM | SM 12:0 | 0.5 |
| PC | PC 13:0_13:0 | 0.5 |
| PC(O) | PC 13:0_13:0 | 0.5 |
| PC(P) | PC 13:0_13:0 | 0.5 |
| LPC | LPC 13:0 | 0.5 |
| LPC(O) | LPC 13:0 | 0.5 |
| PE | PE 17:0_17:0 | 0.5 |
| PE(O) | PE 17:0_17:0 | 0.5 |
| PE(P) | PE 17:0_17:0 | 0.5 |
| LPE | LPE 14:0 | 0.5 |
| PI | PE 17:0_17:0 | 0.5 |
| PS | PS 17:0_17:0 | 0.5 |
| PG | PG 17:0_17:0 | 0.5 |
| CE | CE 18:0 (*d6*) | 20 |
| DG | DG 15:0_15:0 | 5 |
| TG | TG 17:0_17:0_17:0 | 5 |
| FA | FA 15:0 (*d3*) | 0.5 |

#### 2.2.2 Quality assurance (QA) and quality control (QC) measures

1. Two quality control (QC) samples, namely pooled quality control sample (PQC), and a technical quality control sample (TQC, sample with all standards used).
2. Equal amounts from all the samples are pooled together to make the PQC sample.
3. To stabilize the LC/MS system, ten PQC samples are injected prior to injecting experimental samples.
4. PQC samples are injected at equal time intervals to monitor the analytical variability.
5. The lipid extracts without the matrix (here, cell homogenate) are called as TQC samples. They are evenly injected throughout the sample run to ensure the stability of the analytical platform by monitoring the variation in signal intensity over time.
6. Blank extracts are also run at the beginning of sample batch to filter out background ions. All solvents should be HPLC grade.

#### 2.2.3 Lipid separation

1. Reverse-phase liquid chromatography using Shimazu Prominence HPLC system.
2. HPLC column used: Zorbax C18, 1.8 μm, 50 × 2.1 mm column.
3. Flow rate: 300 μL/min using a linear gradient of mobile phase A and mobile phase B.
4. Mobile phase A and B in 10 mM ammonium formate: tetrahydrofuran, methanol, and water in the ratio 20:20:60 (v/v/v) and 75:20:5 (v/v/v), respectively.
5. Elution: begin with 0% solvent B; elevate to 100% B at 8.00 min; keep at 100% B for 2.5 min; and return to 0% B over 0.5 min.

#### 2.2.4 Mass spectral analysis

1. The AbSciex 4000 QTRAP triple quadrupole linear hybrid mass spectrometer is used to introduce lipids eluted from the HPLC system. Operated in scheduled Multiple Reaction Monitoring (MRM) mode.
2. All lipid species are scanned in positive electrospray ionization mode [ESI^+^] mode, except for fatty acids. Instrument settings optimized as: curtain gas (psi), 26; collision gas (nitrogen), medium; ion spray voltage (V), 5500; temperature (°C), 500.0; ion source gas 1 (psi), 40.0; ion and source gas 2 (psi), 30.0.
3. Flow injection analysis is used to fix collision energy and declustering potential for each lipid class. The MRM detection window is fixed between 45 and 90s depending upon the chromatographic peak width of the lipid class.

### 2.3 Lipidomics data analysis

#### 2.3.1 Metaboanalyst v5.0

1. User-friendly, web-based interface metabolite data analysis platform.
2. Used to perform statistical and machine learning analysis, including univariate - t-test, volcano plot, ANOVA; multivariate - principal component analysis (PCA), partial least squares-discriminant analysis (PLS-DA); clustering - dendrogram, and heatmap^30–32^.
3. URL: https://www.metaboanalyst.ca/.

#### 2.3.2 Human Metabolome Database v5.0 (HDMB)

1. It is a free electronic database sourcing detailed information on small molecule metabolites from the human body^33,34^.
2. This database contains or links chemical, clinical, and molecular biology or biochemistry data.
3. There are 220,945 metabolite entries up to November 2023, including water- and lipid-soluble metabolites. Furthermore, 8,610 protein sequences are connected to these metabolite entries.
4. Lastly, HMDB provides unique IDs for each metabolite, such as the SMILES code.
5. URL: https://hmdb.ca/.

#### 2.3.3 Swiss Target Prediction

1. It is a free web-based platform which allows the estimation of the most probable molecular targets of another molecule^35^.
2. The estimation is possible based on 2D and 3D similarities with a library of 370,000 known actives for over 3,000 proteins from 3 distinct species^36^.
3. Given a lipid SMILES code, Swiss Target Prediction can predict possible interacting proteins with that metabolite (SMILES).
4. URL: http://www.swisstargetprediction.ch/.

#### 2.3.4 String v12.0

1. This is a user-friendly free database of known and predicted protein-protein interactions, either physical (direct) or functional (indirect) associations^37^.
2. The associations are originated from computational prediction coming from knowledge transfer and combination with other databases^38,39^.
3. URL: https://string-db.org/.

## 3 Methods

### 3.1 A549 cell culture

1. Carry out cell culture in biosafety cabinet in sterile condition.
2. Culture A549 lung adenocarcinoma cells in both adherent and non-adherent plates (100mm); the former mimics primary tumor conditions (attached) and the latter metastatic tumors in circulation (detached). Use high glucose growth media. Three plates of cells for each condition *(see* ***Note 1***).
3. Grow cells until reach 100% confluence, maintained in a humidified incubator with 95% air and 5% CO_2_ at 37 °C. Total yield per plate is approximately 5 million cells.
4. Once reached 100% confluence, collect cells using 1x PBS, in a 1.5mL tube.
5. Sonicate samples (3 cycles, 5 seconds each).
6. Centrifuge samples (1000g for 10 minutes).
7. Collect the supernatant for each sample and transfer to a newly labeled tube.
8. Measure protein concentration using Lowry Protein Assay method.
9. Ideally 1mg/mL of sample is necessary for lipidomics. Therefore, normalize all proteins concentration to 1mg/mL.
10. Proceed to lipidomics sample preparation and lipidomics analysis *(see* ***Note 2***).

### 3.2 Lipid chromatography and mass spectrometry^29^

#### 3.2.1 Lipid Expression

1. Lipid extraction was performed using chloroform and methanol^29,40^.
2. 2. In a new 1.5 mL tube polypropylene tube, add 100 μL of cell homogenate, and 30 μL of the internal standard (ISTD) plus 200 μL of chloroform/methanol (2:1, v/v) creating a new mixture *(see* ***Note 3***).
3. Vortex mixture on a rotary mixer for 10 min and sonicated in a water bath at room temperature (RT) for 30 min.
4. Allow mixture to settle for 20 min and then centrifuged (20,000g, 20 min, RT).
5. Transfer the upper lipid-containing phase into a clean polypropylene tube and dry under a stream of nitrogen gas at RT.
6. Reconstitute lipids in 50 μL water-saturated 1-butanol and sonicate for 10 min (water bath, RT).
7. Add 50 μL of 10 mM ammonium formate in methanol to final lipid extract.
8. Centrifuge the extract (10,000g, 10 min, RT).
9. Transfer 80 μL of the supernatant into the micro insert in sample vials for lipid analysis.

#### 3.2.2 Lipid separation and chromatography

1. Separate lipids on a reverse-phase liquid chromatography-electrospray ionization tandem mass spectrometry (LC-ESI-MS/MS) platform using a Prominence chromatographic system.
2. Use Analyst v1.6 and Multi-Quant v2.1 software for instrument control and data processing.
3. Perform lipid separation on a Zorbax C18, 1.8 μm, 50 × 2.1 mm column at a flow rate set to 300 μL/min using a linear gradient of mobile phase A and mobile phase B *(see* ***Note 4***).
4. Re-equilibrate the column to the starting conditions (0% mobile phase B) for 3 min prior the next sample injection.

#### 3.2.3 Mass spectral analysis

1. Add to AbSciex 4000 QTRAP triple quadrupole linear hybrid mass spectrometer the eluted lipids from HPLC.
2. Mass spectrometer operation schedule mode: Multiple Reaction Monitoring (MRM).
3. Screen total of 322 unique lipids, 25 different lipid classes/subclasses for targeted semi-quantitation (**Supplementary Table S3**, Reference 29).
4. Scan all lipid species (except fatty acids) positive electrospray ionization mode [ESI^+^] mode.
5. Identify the individual lipids in each lipid class by using lipid class-specific precursor ions or neutral losses (**Supplementary Table S2**, Reference 29).
6. Lipids are represented by the total carbon number of the fatty acids.
7. In the ESI^+^ mode, optimize the instrument settings as follows: curtain gas (psi), 26; collision gas (nitrogen), medium; ion spray voltage (V), 5500; temperature (°C), 500.0; ion source gas 1 (psi), 40.0; ion and source gas 2 (psi), 30.0.
8. Fix the MRM detection window between 45 and 90s depending upon the chromatographic peak width of the lipid class.
9. Scan M^+1^/M^+2^ isotopes for high-intensity lipid species to prevent detection saturation.
10. Proceed to data analysis once lipid abundance has been detected.

### 3.3 Lipidomics data analysis

#### 3.1.1 Metaboanalyst 5.0

1. Create a matrix excel table with all lipids and labels and export to CSV format then upload it to Metaboanalyst (see ***Note 5***).
2. Normalize lipid abundance of the samples to 1mg/mL to match protein determination normalization parameters.
3. Next, ensure there is no empty values (see ***Note 6***).
4. First column should be listed all lipids, and following columns should be listed all groups, including a minimum of 3 replicates per group.
5. Add two rows and name “A1” as “Row Labels, followed by group labels (A549 A_1, A549 A_2, A549 A_3, A549 D_1, A549 D_2, A549 D_3). On row”A2”, name as “Labels”, followed by group labels without replicate number (A549 Attached, A549 Attached, A549 Attached, A549 Detached, A549 Detached, A549 Detached), as shown in **Figure 1**.
6. Once label formatting is done, export excel table as comma delimited format (CSV).
7. Open Metaboanalyst website.
8. Click on red button: “click here to start”.
9. A new page will be opened with “module overview” of all available functions. Click on “Statistical Analysis [1 Factor]”, under impute data type “generic format .csv or .txt table files”.
10. Next, upload data under the option “A plain text file (.txt or .csv)”. And select the following options: Data Type “Concentrations”; Format “Samples in columns, unpaired”; Data File: “Choose” – upload csv formatted matrix/table.
11. “Data integrity check” page will be prompted. If matrix was well formatted, at this stage it will indicate the number of groups, samples and compounds identified. Click on “Proceed”.
12. Data Filtering: under “Statistical Filters”, select “mean intensity value”, then “Proceed”.
13. A “Normalization overview” page will be prompted, select the following options: “Sample normalization – none”, “Data transformation – none”, “Data scaling – auto scaling”. Click on “Normalize”, next “View Results”, next “Proceed”.
14. Next, under “Select an analysis path to explore” new page you can choose type of analysis to run and generate graphics.
15. Metaboanalyst results *(see* ***Note 7***).
16. On “Select an analysis path to explore” page, under drop down menu located on the left side of the page, click on “Statistics”.
17. Next, click on “Heatmap” (**Figure 2**) to visualize overview lipid abundance and select the following settings:
  1. Data source: normalized data.
  2. Standardization: autoscale features.
  3. Distance measure: eucledian.
  4. Clustering: ward.
  5. Color contrast: default.
  6. Font size: 10.
  7. View mode: overview.
  8. Other view options: show heatmap color legend, samples/T-test ANOVA.
  9. To visualize most significant lipid abundance and lipid names, under “Other view options”, select “Use top”, then enter the number of lipids to be displayed, e.g. 50, as shown in **Figure 3**.
  10. Next, click “Show row names”.
18. Back to “Statistics” menu, select “Volcano Plot” (see **Note 8**).
19. Select the following settings to generate Volcano Plot graph (**Figure 4**):
  1. Analysis: unpaired.
  2. Plot style: show label.
  3. Theme: blackwhite.
  4. X-axis: Fold Change threshold 2.0; Direction of comparison: A5459 Detached/A549 Attached.
  5. Y-axis: p-Value threshold FDR 0.05; group variance equal *(see* ***Note 9***).
20. Back to “Statistics” menu, select “PCA” for unsupervised analysis (**Figure 5**).
21. A new page “Principal Component Analysis (PCA)” will be prompted. Choose the following parameters:
  1. Select “2D Scores Plot.
  2. Specify PC on X-axis: 1.
  3. Specify PC on Y-axis: 2
  4. Display 95% confidence regions: check.
  5. Display sample names: check.
22. Back to “Statistics” menu, select “PLSDA”, supervised analysis (**Figure 6**).
  1. A new page “Partial Least Squares Discriminant Analysis (PLS-DA)” will be prompted.
  2. Under 2D scores plot, apply same settings as indicated for “PCA”
23. Still within PLSDA menu, click on “Imp. Features” to measure the PLS-DA variable importance in projection (VIP) score (**Figure 7**), and apply the following settings:
  1. Importance measure: VIP Score, comp 1.
  2. Show top features (display on graph should be min 1.2 VIP score).
  3. Download excel sheet with all lipids with VIP score >1.4 and proceed to next steps of analysis to predict lipid-protein interaction.

**Figure 1.**
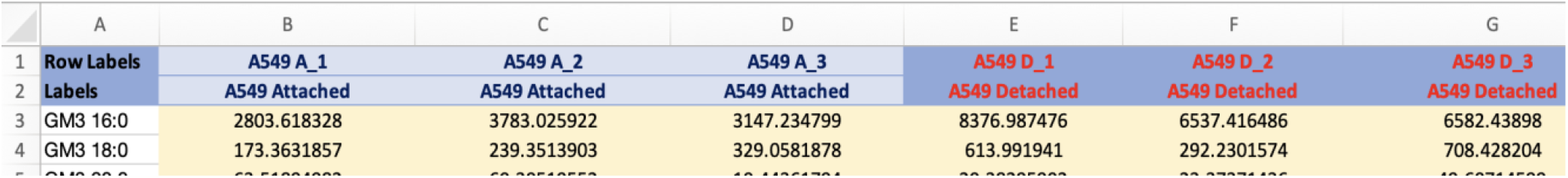
Screenshot of matrix formatting style to be uploaded at Metaboanalyst.

**Figure 2.**
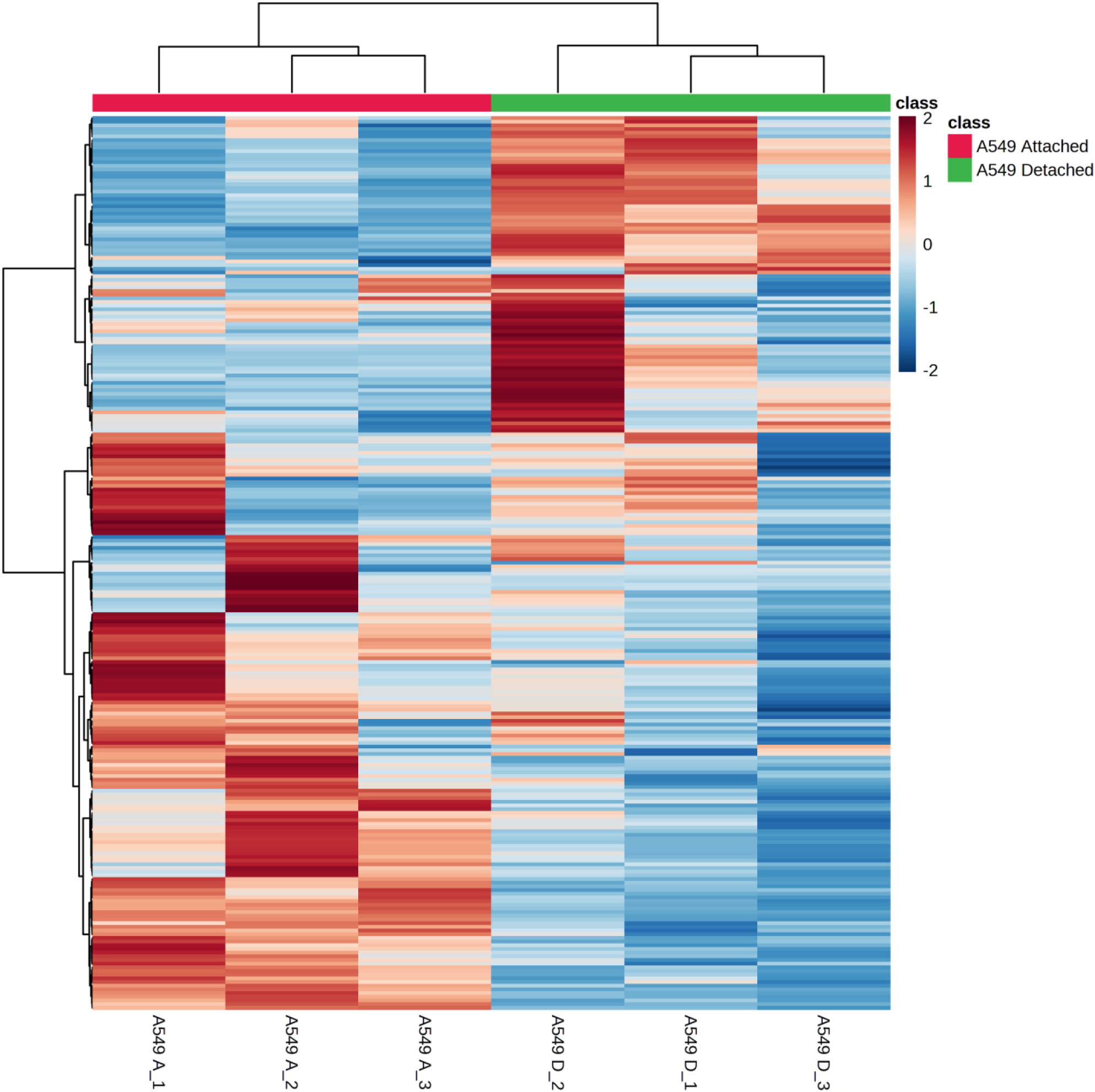
Heatmap of all lipids abundance in A549 attached and A549 detached conditions. Generated in Metaboanalyst.

**Figure 3.**
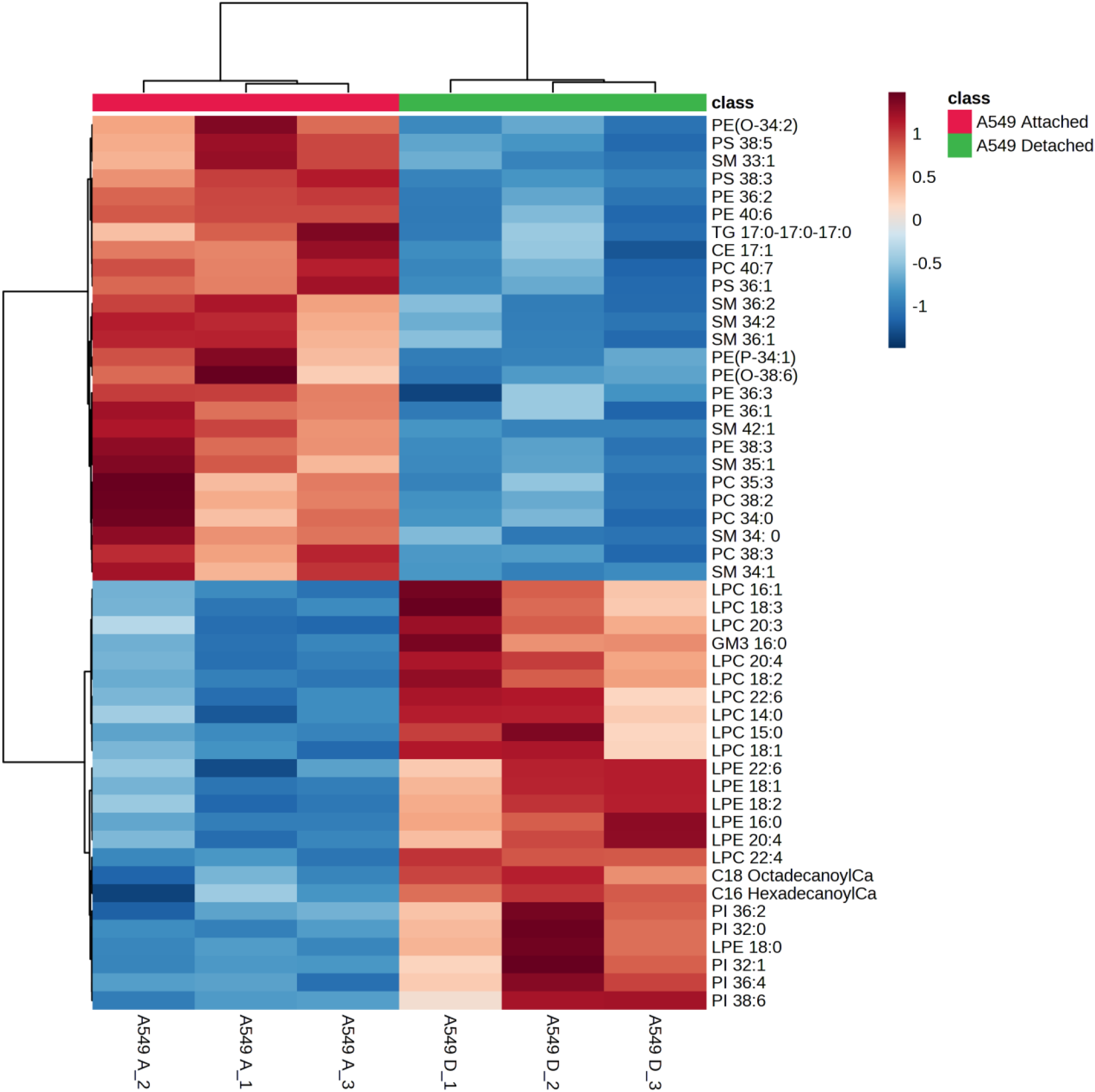
Heatmap of top 50 lipids abundance in A549 attached and A549 detached conditions. Generated in Metaboanalyst.

**Figure 4.**
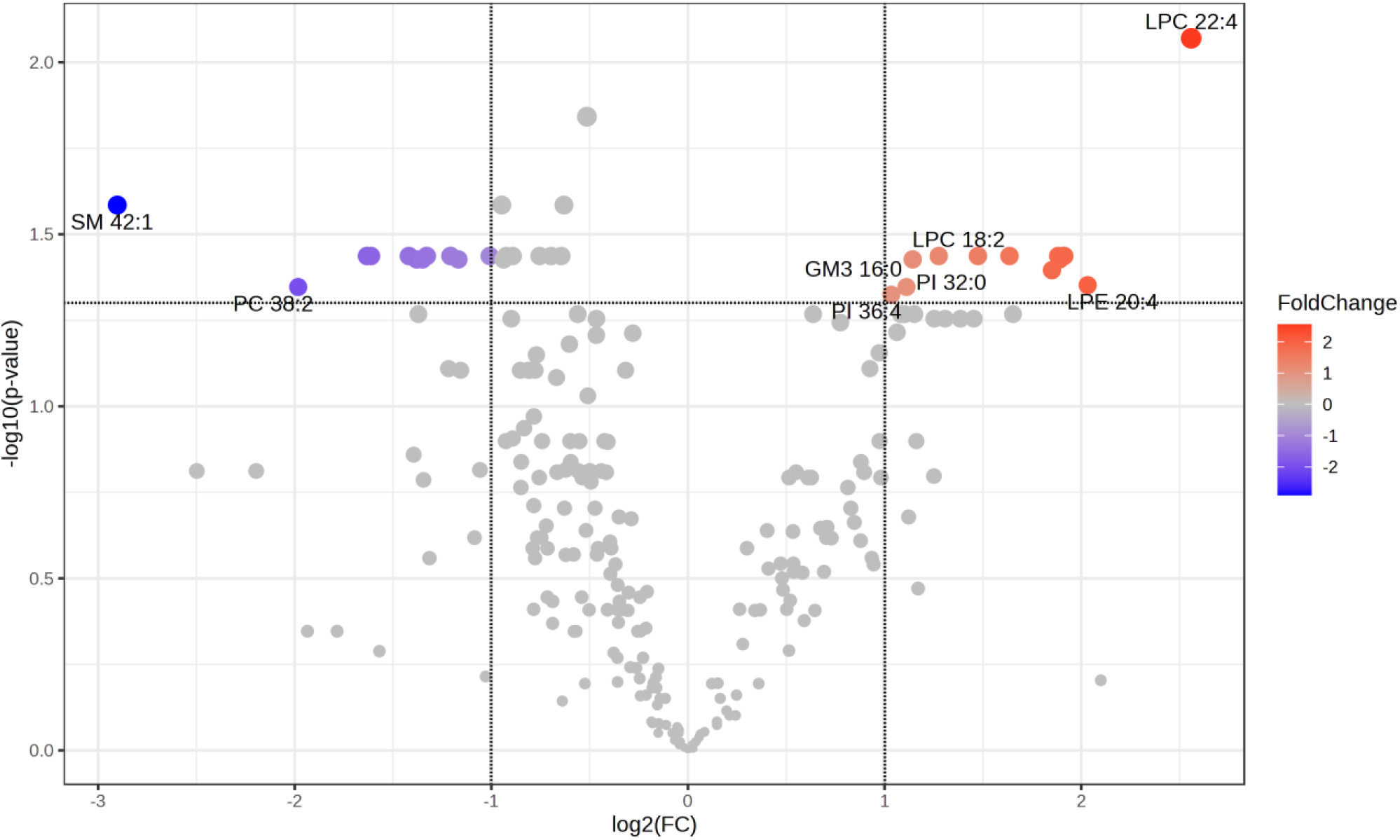
Volcano plot of A549 detached condition versus A549 attached condition, displaying the most significant lipids either upregulated (red) or downregulated (blue), based on FC 2 and FDR 0.05.

**Figure 5.**
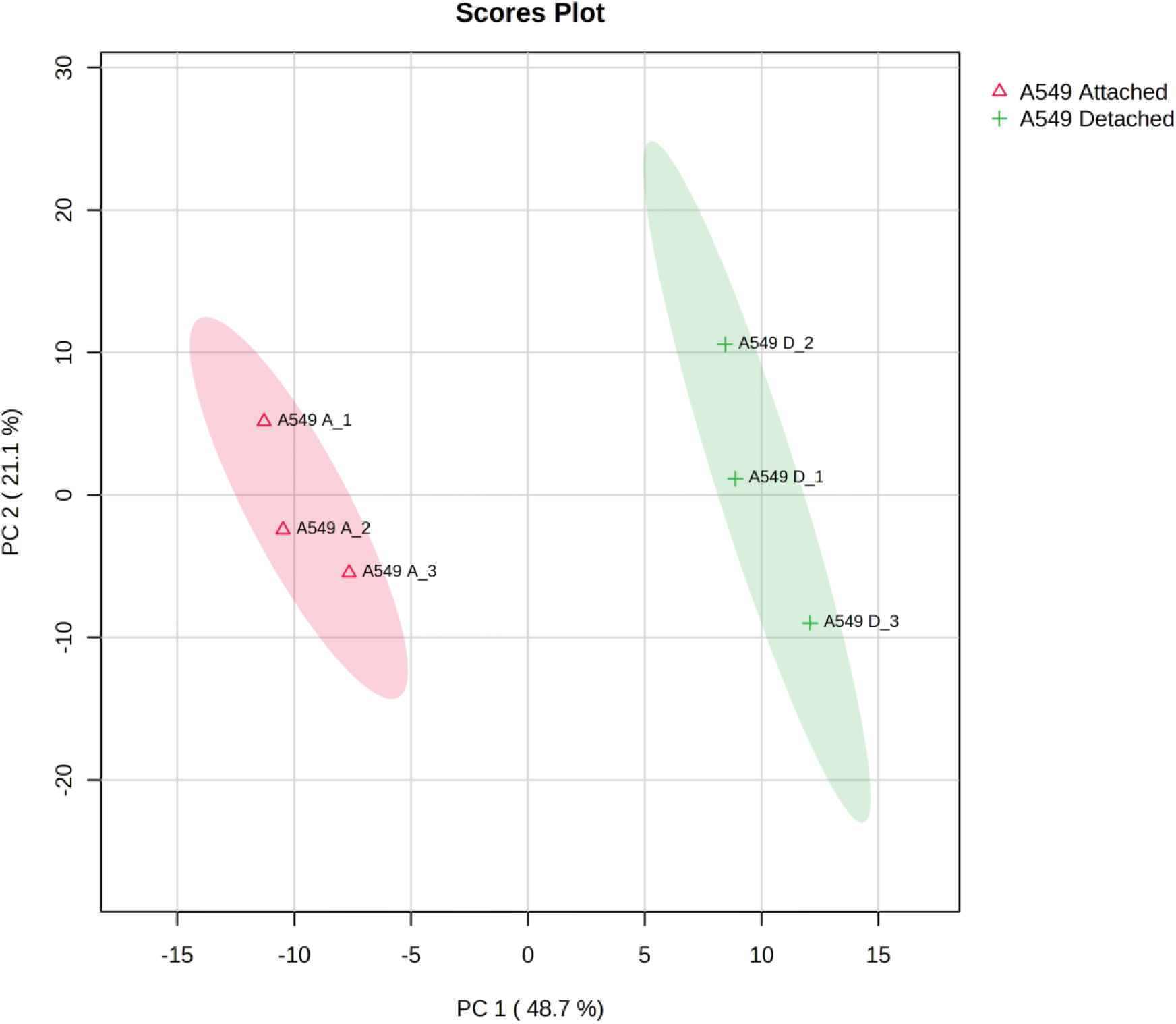
Principal Component Analysis (PCA) plot of A549 attached and A549 detached conditions, unsupervised analysis, showing distinct biological variance between groups.

**Figure 6.**
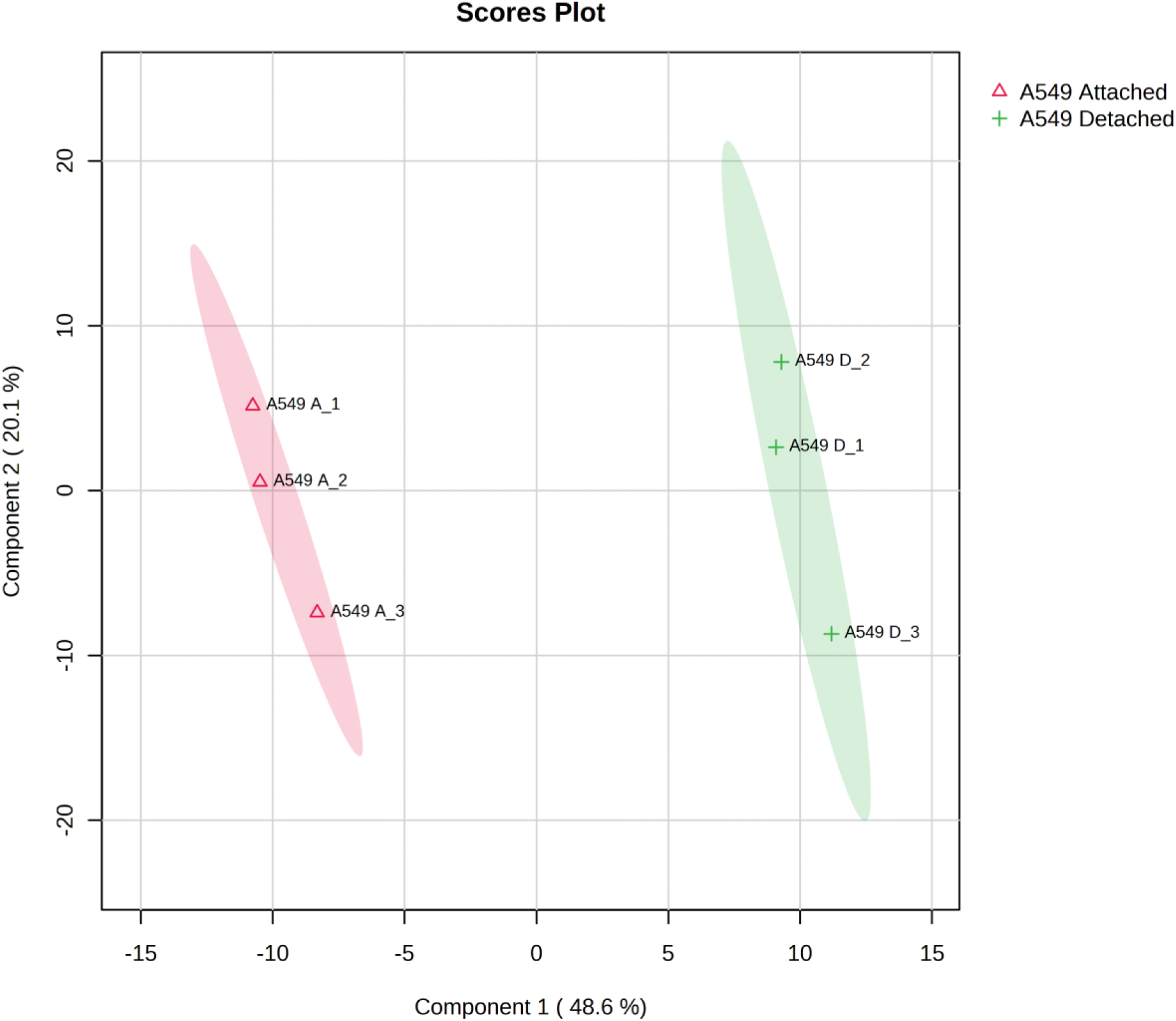
Partial Square Least Discriminant Analysis (PLSDA) plot of A549 attached and A549 detached conditions, supervised analysis, showing distinct biological variance between groups.

**Figure 7.**
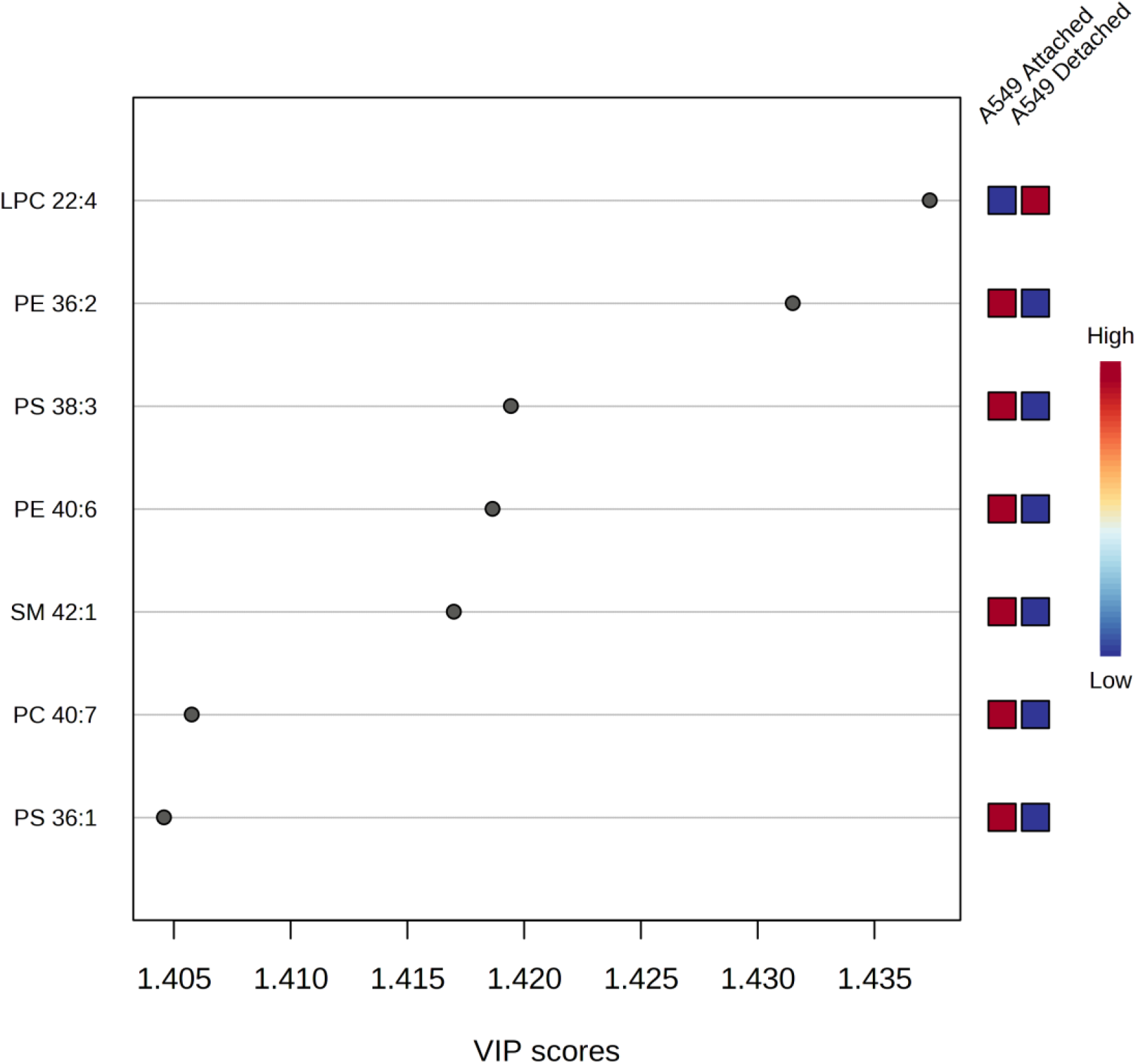
Partial Square Least Discriminant Analysis (PLSDA) variable importance in projection (VIP) score plot of A549 attached and A549 detached conditions (VIP >1.4).

#### 3.3.2 Human Metabolome Database (HMDB)

1. At this step, all significant lipids selected from VIP score >1.4 will be searched at HMDB to identity their unique ID and smiles code. This smiles code will be used to predict possible interacting proteins with each metabolite.
2. First, create a new excel table with PC1 exported from VIP score tab on Metaboanalyst (**Figure 8)**. In front of each lipid, add columns for corresponding VIP-PC1 value, lipid common name, and smiles code.
3. Enter lipid name in HMDB search engine and look for exact lipid. Click on the correct lipid and copy “lipid common name” and “smiles code” and past to excel table. Repeat this step for all significant lipids (VIP >1.4 in this case).
4. Proceed to Swiss Target Prediction to identify interacting proteins with each metabolite/lipid in query.

**Figure 8.**
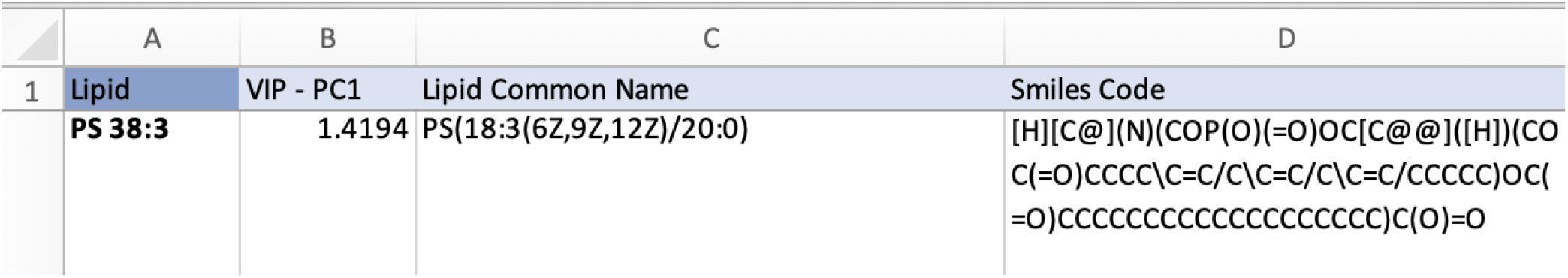
Screenshot of template table to annotate significant lipids details, including VIP score >1.4, lipid common name and smiles code obtained from HMDB.

#### 3.3.3 Swiss Target Prediction

1. First, add a new column to excel sheet, and label as “Uniprot ID”, where top 15 predicted interacting proteins will be pasted (**Figure 9)**.
2. On the Swiss Target Prediction main webpage, follow the following steps:
  1. Select a species: homo sapiens.
  2. Paste a smile code in the allocated box.
  3. Click on red button “Predict targets”, it may take a few minutes to load.
  4. A new page will be prompted with a list of predicted interacting proteins.
  5. Click under “show entries” and select 15.
  6. Lastly, under “Export Results” tab, select icon “CSV” and choose folder location for download.
  7. Repeat this step for all lipids in query.
  8. After all saved all CSV files containing predicted interacting proteins for all lipids, now open each one individually, and copy the protein ID of top 15 targets, and past in the new column created in excel sheet (within the same cell), labeled “Uniprot ID”.
  9. Once excel table is complete with all corresponding protein targets, proceed to pathway enrichment analysis using String.

**Figure 9.**
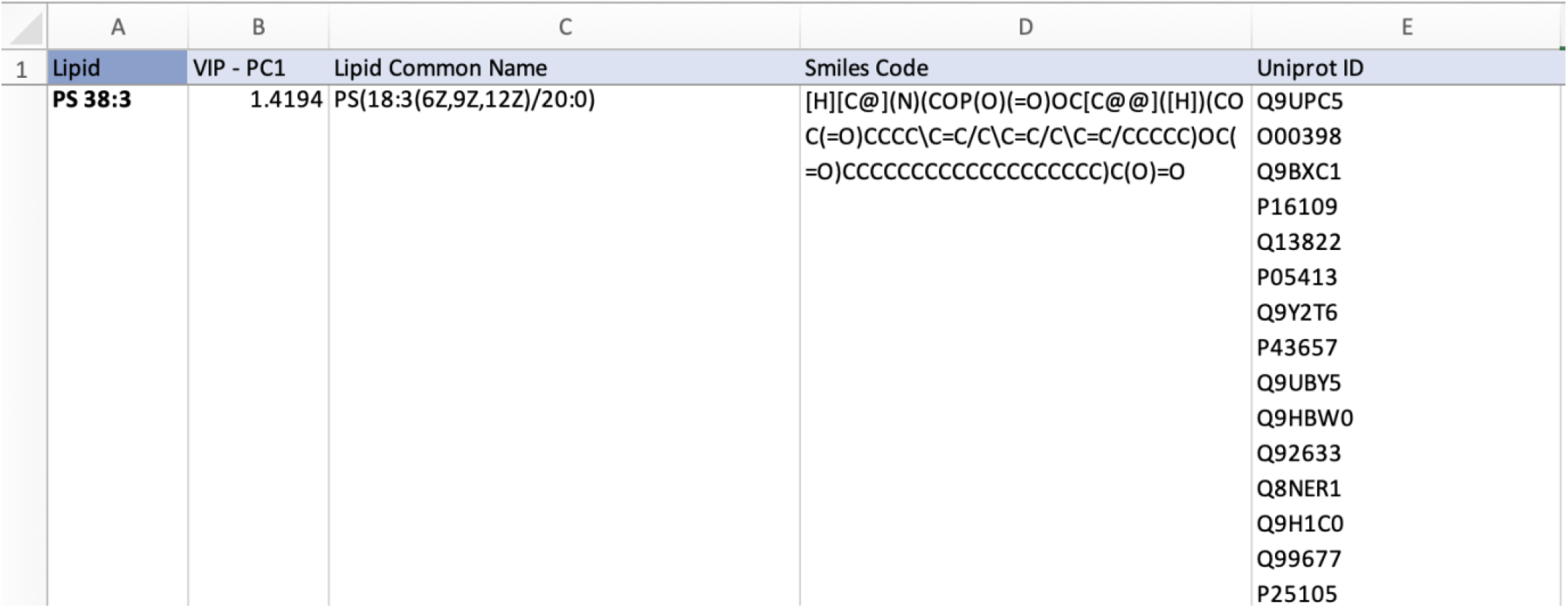
Screenshot of template table to annotate significant lipids details, including VIP score >1.4, lipid common name, smiles code obtained from HMDB, and Uniprot IDs obtained from SwissTargetPrediction.

#### 3.3.4 String

1. On the String main webpage, follow the following steps:
  1. Left menu tab: click on “Multiple proteins”.
  2. Paste list of all protein ID from excel sheet, in a column, one protein per line *(see* ***Note 10***).
  3. Select organism: *Homo sapiens*, and click “search”.
  4. A list of matches will prompt, click “continue”.
  5. The network will be now generated, and a new panel of commands are available to adjust and set statistical parameters for the analysis.
  6. Click on “Settings” and set the following parameters for “Basic Settings”:
    1. Network type: full string network.
    2. Meaning of network edges: confidence.
    3. Active interaction sources: check all.
    4. Minimum required interaction score: high confidence 0.700.
    5. Max number of interactions to show: 1^st^ shell (none/query proteins only).
  7. Still under “Settings”, set the following parameters for “Advanced Settings”:
    1. Network display mode: interactive.
    2. Network display options: check boxes for “enable 3D bubble design”, “disable structure previews inside network bubbles”, “center protein names on nodes”, “protein font size 14”.
  8. Click on “Update” to save new settings.
  9. Next, click on “Analysis” tab.
  10. Look for “KEGG” pathways, then click on “FDR” so pathways will be listed based on most significant FDR values, with arrow pointing up.
  11. Click on KEGG pathway of interest listed, such as the “PI3K-AKT signaling pathway” (FDR 5.44e-05). It will be highlighted in the network the proteins from your query involved in this pathway, shown in red (**Figure 10**).
  12. Still under “Analysis” tab, go to the bottom of the page and click on “download” “KEGG pathways” (**Table 2)**.
  13. Next, click on “Exports” tab.
  14. Under “Export your current network”, download following files “high-resolution bitmap” network image, and “functional annotations” table containing all analysis results, including KEGG pathways.

**Figure 10.**
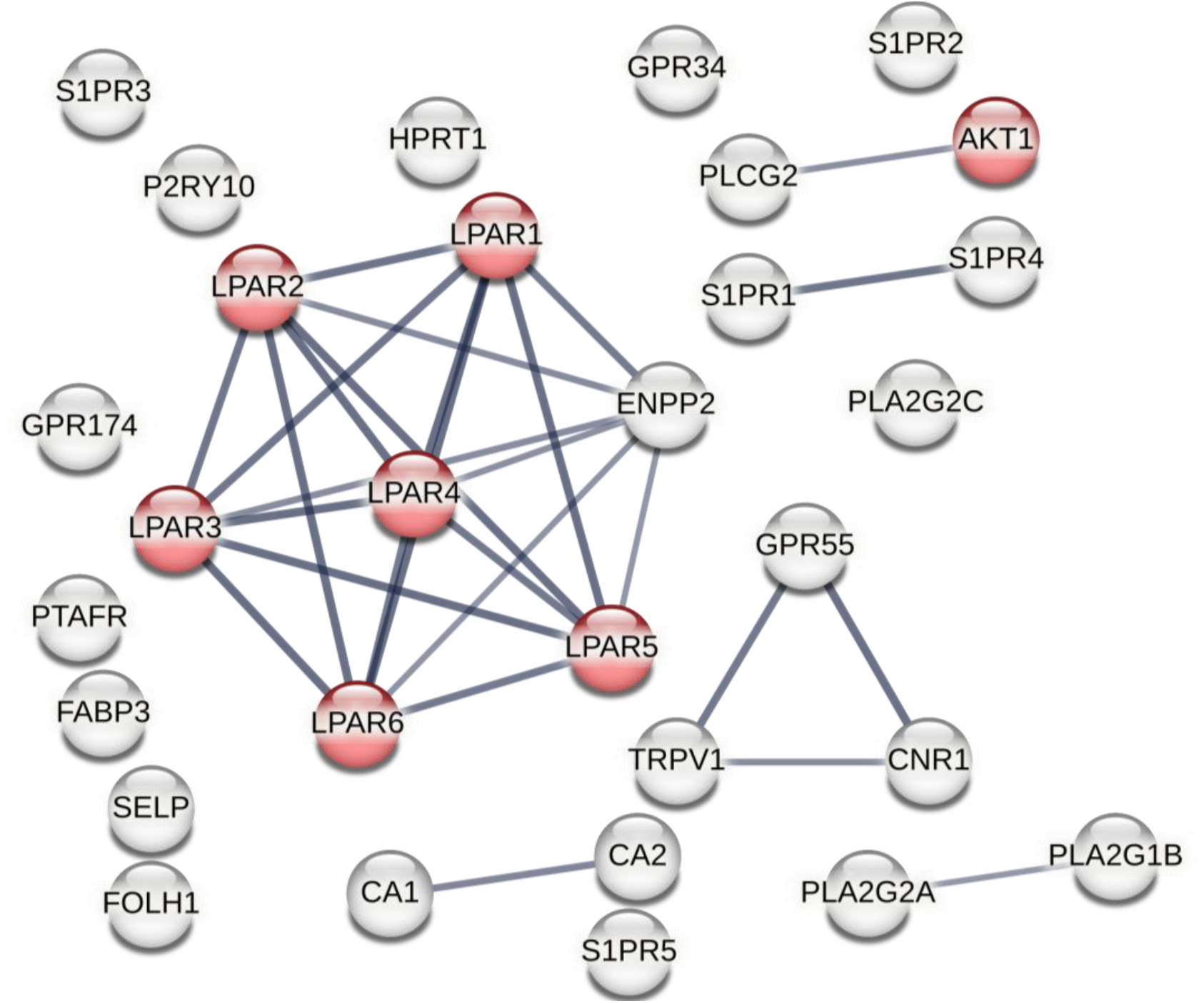
String network analysis highlighting PI3K-AKT signaling pathway (KEGG) in red nodes (FDR 5.44e-05). Network was generated based on significant lipids VIP >1.4 and respective predicted interacting proteins. Minimum 15 possible interacting proteins were entered on String, per lipid. Confidence interaction min 0.700.

**Table 2.** List of significant enriched KEGG pathways, highlighting PI3K-Akt signaling as one of the main correlation pathways to autophagy changes in A549 adenocarcinoma primary and metastatic tumors. KEGG) in red nodes (FDR 5.44e-05). Network was generated based on significant lipids VIP >1.4 and respective predicted interacting proteins. Minimum 15 possible interacting proteins were entered on String, per lipid.

| #term ID | term description | observed gene count | background gene count | strength | false discovery rate |
| --- | --- | --- | --- | --- | --- |
| hsa04080 | Neuroactive ligand-receptor interaction | 14 | 329 | 1.45 | 6.73E-15 |
| hsa04072 | Phospholipase D signaling pathway | 8 | 147 | 1.55 | 1.03E-08 |
| hsa04015 | Rap1 signaling pathway | 7 | 201 | 1.36 | 2.43E-06 |
| hsa04071 | Sphingolipid signaling pathway | 6 | 116 | 1.53 | 2.43E-06 |
| hsa04151 | PI3K-Akt signaling pathway | 7 | 349 | 1.12 | 5.44E-05 |
| hsa05200 | Pathways in cancer | 8 | 515 | 1.01 | 5.44E-05 |
| hsa00592 | alpha-Linolenic acid metabolism | 3 | 25 | 1.9 | 0.00049 |
| hsa00591 | Linoleic acid metabolism | 3 | 29 | 1.83 | 0.00064 |
| hsa04972 | Pancreatic secretion | 4 | 97 | 1.43 | 0.00064 |
| hsa04014 | Ras signaling pathway | 5 | 225 | 1.16 | 0.00078 |
| hsa04975 | Fat digestion and absorption | 3 | 41 | 1.68 | 0.0012 |
| hsa00565 | Ether lipid metabolism | 3 | 47 | 1.62 | 0.0017 |
| hsa00590 | Arachidonic acid metabolism | 3 | 60 | 1.52 | 0.0031 |
| hsa05130 | Pathogenic Escherichia coli infection | 4 | 187 | 1.15 | 0.0046 |
| hsa04810 | Regulation of actin cytoskeleton | 4 | 209 | 1.1 | 0.0065 |
| hsa00910 | Nitrogen metabolism | 2 | 17 | 1.89 | 0.0079 |
| hsa00564 | Glycerophospholipid metabolism | 3 | 97 | 1.31 | 0.0092 |
| hsa04068 | FoxO signaling pathway | 3 | 126 | 1.19 | 0.0182 |
| hsa01100 | Metabolic pathways | 8 | 1435 | 0.56 | 0.0194 |
| hsa04270 | Vascular smooth muscle contraction | 3 | 132 | 1.17 | 0.0194 |

## 4 Conclusion

Our results showed a distinct biological variance among A549 adenocarcinoma attached and detached cells. Based on Metaboanlyst lipidomics analysis, we could observe a significant increase in lysophosphatidylcholines (LPC), phosphatidylcholines (PC), phosphatidylethanolamine (PE), phosphatidylserine (PS), phosphatidylethanolamine (PE), and sphingomyelin (SM) lipids in A549 attached cell group, while decreased in detached group. We further predicted the possible interacting proteins for each significant lipid and performed an enrichment analysis based on the Kyoto Encyclopedia of Genes and Genome (KEGG). The KEGG results indicated that the PI3K-AKT pathway is among the most significant enriched pathways. Furthermore, PI3K-AKT are at the core of mTOR and autophagy signaling pathways regulation. Hence, this method may serve as an advanced approach for exploratory research of the role of lipids in autophagy and cancer, and further experimental data analysis are necessary to validate the predicted biological findings *(see* ***Note 11***).

## 5 Notes

1. **Note 1:** Make sure to have minimum triplicates of each condition. Ideally, best if four or more replicates of each group/condition. The main reason is ensuring repeatability, consistency, and eliminate any variability or possible outliers. Moreover, a minimum of three replicates is required to perform statistical analysis. Also, make sure cells are serum starved and grown in 1% ITS as indicated in material section. The reason is because FBS may interfere in the circulating levels of chemokines and cytokins, while ITS does not impact it.
2. **Note 2:** A typical workflow of lipidomics analysis of biological samples consists of sample collection and storage, homogenization, internal standard addition for calibration, analysis of lipids with and without separation, data processing and lipid identification, and biomedical application.
3. **Note 3:** Lipid internal standards are either odd-chain or deuterated and not present endogenously.
4. **Note 4:** Maintain analytical column and autosampler at 50°C and 25°C, respectively, during analysis. The injection volume is 5 μL.
5. **Note 5:** All lipids should be abbreviated with character free. Ensure that all text is exactly the same within replicates nomenclature. If a tiny space is added extra in one replicate, it will be indicated as a new group and will be misinterpreted by Metaboanalyst, which is an R-based web analysis tool.
6. **Note 6:** It is important to note that this procedure does not account for all missing values, which must be eliminated for subsequent data analysis procedures such as principal component analysis (PCA). Rather than relying solely on the imputation of zero values, it is advantageous to use data-driven imputation methods such as Random Forest (RF), k-nearest neighbors (KNN) and Singular Value Decomposition (SVD) or use the Limit of Detection (LOD) value. The sample size generally had only a minimal influence on the results. However, with a sample size of N = 10, even a moderate amount of missing data significantly reduced the statistical power. In cases where more data is missing, KNN and RF prove to be effective means of statistical testing, causing minimal delta and standard deviation (SD) bias^41^.
7. **Note 7:** All generated graphs in Metaboanalyst can be saved by clicking on a “paint palette” icon, usually displayed besides each graphic. When available, “Excel” icons will be displayed, indicating that a table is available for download.
8. **Note 8:** Volcano plots are generated with two group comparison at a time. If your matrix contains 3 groups or more, you will have to select only two groups at a time to perform analysis. Groups can be edited under left menu tab on metaboanlyst by clicking on “Data processing”, then select “Data Editor”, manually select chosen groups, and start data normalization steps once again.
9. **Note 9:** Ideally, false discovery rates (FDR) should be set between 0.01 to 0.05 for the identification of significant statistical changes.
10. **Note 10:** Make sure no special character is added before or after any proteins. Sometimes it may occur to automatically be added quotations (“ “) when pasting from excel. Quickly scan over the list of proteins and remove any special characters that have may been automatically added. If this step is not done, there will be errors.
11. **Note 11:** The lipid-protein interaction is a bioinformatic prediction approach used, which may not be as accurate as if performed a sample proteomics analysis. However, in the absence of proteomics cellular data, the in-silico screening is a great start and indication of possible target proteins to be considered.

